# The sex-specific VC neurons are mechanically activated motor neurons that facilitate serotonin-induced egg laying in *C. elegans*

**DOI:** 10.1101/2020.08.11.246942

**Authors:** Richard J. Kopchock, Bhavya Ravi, Addys Bode, Kevin M. Collins

## Abstract

Successful execution of behavior requires coordinated activity and communication between multiple cell types. Studies using the relatively simple neural circuits of invertebrates have helped to uncover how conserved molecular and cellular signaling events shape animal behavior. To understand the mechanisms underlying neural circuit activity and behavior, we have been studying a simple circuit that drives egg-laying behavior in the nematode worm *C. elegans*. Here we show that the sex-specific, Ventral C (VC) motor neurons are important for vulval muscle contractility and egg laying in response to serotonin. Ca^2+^ imaging experiments show the VCs are active during times of vulval muscle contraction and vulval opening, and optogenetic stimulation of the VCs promotes vulval muscle Ca^2+^ activity. Blocking VC neurotransmission inhibits egg laying in response to serotonin and increases the failure rate of egg-laying attempts, indicating that VC signaling facilitates full vulval muscle contraction and opening of the vulva for efficient egg laying. We also find the VCs are mechanically activated in response to vulval opening. Optogenetic stimulation of the vulval muscles is sufficient to drive VC Ca^2+^ activity and requires muscle contractility, showing the presynaptic VCs and the postsynaptic vulval muscles can mutually excite each other. Together, our results demonstrate that the VC neurons facilitate efficient execution of egg-laying behavior by coordinating postsynaptic muscle contractility in response to serotonin and mechanosensory feedback.

**Significance Statement:** Many animal motor behaviors are modulated by the neurotransmitters serotonin and acetylcholine. Such motor circuits also respond to mechanosensory feedback, but how neurotransmitters and mechanoreceptors work together to coordinate behavior is not well understood. We address these questions using the egg-laying circuit in *C. elegans* where we can manipulate presynaptic neuron and postsynaptic muscle activity in behaving animals while recording circuit responses through Ca^2+^ imaging. We find that the cholinergic VC motoneurons are important for proper vulval muscle contractility and egg laying in response to serotonin. Muscle contraction also activates the VCs, forming a positive feedback loop that promotes full contraction for egg release. In all, mechanosensory feedback provides a parallel form of modulation that shapes circuit responses to neurotransmitters.

## Introduction

A fundamental goal of neuroscience is to understand the neural basis of behavior (Bargmann & Marder, 2013). Recent work reporting the synaptic wiring diagrams, or connectomes, of nervous systems provides an unprecedented opportunity to study how the nervous system directs animal behavior (Cook et al., 2019; Lerner et al., 2016; Meinertzhagen, 2018). However, connectomes alone are not sufficient to predict nervous system function (Batista-García-Ramó & Fernández-Verdecia, 2018; Swanson & Lichtman, 2016; Taylor et al., 2019). Released neurotransmitters can signal both synaptically and extrasynaptically through distinct receptors to drive short-term and long-term behavior changes (Chase et al., 2004; Del-Bel & De-Miguel, 2018; Donnelly et al., 2013; Hardingham & Bading, 2010; Koelle, 2018). Neuropeptides can be co-released from synapses and signal alongside neurotransmitters (Brewer et al., 2019; Nusbaum et al., 2017). Understanding how an assembly of ionotropic and metabotropic signaling events drives the complex pattern of circuit activity is facilitated by direct tests in invertebrate animals (Bargmann & Marder, 2013) such as those amenable to genetic investigation (Sengupta & Samuel, 2009). Such studies have the potential to reveal conserved neural circuit signaling mechanisms that underlie behavior.

The egg-laying circuit of the nematode worm *C. elegans* provides an ideal model for such a reductionist approach (Figure 1A-B). The canonical egg-laying circuit is comprised of three main cell types: the serotonergic HSN command motor neurons, the vulval muscles, and the cholinergic VC motor neurons (Schafer, 2006). An active egg-laying behavior state is initiated when the HSN neurons release serotonin (Desai et al., 1988; Waggoner et al., 1998), which signals through G protein coupled serotonin receptors (Fernandez et al., 2020; Hapiak et al., 2009), and NLP-3 neuropeptides (Brewer et al., 2019). Individual eggs are laid when the vm1 and vm2 vulval muscle cells contract in synchrony to open the vulva (Collins & Koelle, 2013; Li et al., 2013). The VC motor neurons are the primary cholinergic neurons of the egg-laying circuit (Duerr et al., 2001; Pereira et al., 2015), and are active during egg-laying behavior (Collins et al., 2016; Zhang et al., 2008), but the function of their signaling is not well understood (Schafer, 2006). As in mammals, most muscle contraction events in *C. elegans* are ultimately driven by acetylcholine (ACh; Richmond & Jorgensen, 1999; Trojanowski et al., 2016). Nicotinic ACh receptor (nAChR) agonists promote egg laying by acting on the vulval muscles (Kim et al., 2001; Waggoner et al., 2000), consistent with the VCs and/or other motor neurons releasing ACh to excite the vulval muscles (Waggoner et al., 1998). However, ACh synthesis and packaging mutants are hyperactive for egg laying, indicating ACh can also inhibit egg laying, possibly by signaling through inhibitory muscarinic ACh receptors (Bany et al., 2003; Fernandez et al., 2020). This hyperactive egg-laying phenotype resembles animals in which VC neuron development has been disrupted by laser ablation or mutation (Bany et al., 2003), suggesting instead that ACh released from the VCs acts, at least in part, to inhibit egg laying, perhaps in response to sensory input. The VCs extend processes along the vulva, leading to the proposal they might also relay mechanosensory feedback in response to vulval opening (Zhang et al., 2010). How the VC neurons become activated and signal to regulate egg-laying behavior remains unclear. Such insight is necessary for each cell in a neural circuit to transform static information from wiring diagrams into dynamic and meaningful understanding of animal behavior.

**Figure 1.**
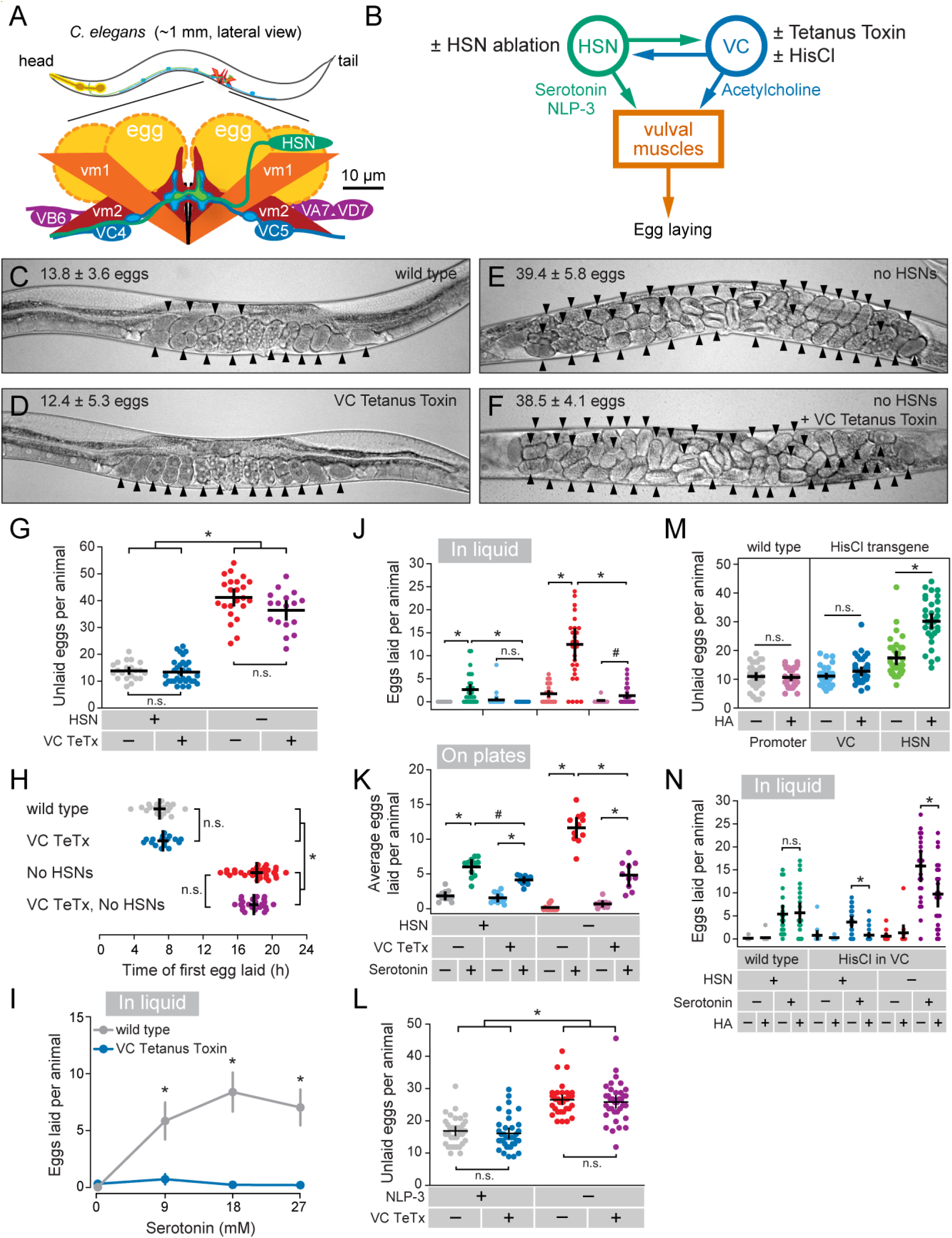
VC neurotransmission facilitates egg laying in response to serotonin. **(A)** Graphical representation of the *C. elegans* egg-laying circuit (modified from Collins et al., 2016). (**B**) Simplified circuit diagram showing the synapses and the primary neurotransmitters released between the HSN neurons, VC neurons, and vulval muscles. The cell-specific transgene expressions performed throughout this figure are indicated. (**C-F**) Representative images of the *C. elegans* uterus showing unlaid eggs (arrowheads) in wild-type, transgenic animals expressing Tetanus Toxin (TeTx) in the VCs, and HSN-ablated *egl-1(986dm)* mutant animals. (**G**) Measurement of steady-state egg accumulation in animals from Figure 1C-F (±95 confidence intervals for the mean); asterisks indicate *p*<0.0001 and n.s. indicates *p*>0.05 (one-way ANOVA with Bonferroni’s correction for multiple comparisons; n≥17). **(H)** Scatter plot showing timing of first egg laid after the larval to adult molt in animals with blocked VC neurotransmission and ablated HSNs (asterisk indicates *p*<0.0001, one-way ANOVA with Bonferroni’s correction for multiple comparisons; n≥17). (**I**) Blockage of VC synaptic transmission inhibits serotonin-induced egg laying. Animals expressing TeTx in the VCs were placed into M9 buffer or M9 containing the indicated concentrations of serotonin and scored for number of eggs laid; asterisks indicate *p*<0.0001 (Kruskal-Wallis test with Dunn’s correction for multiple comparisons; n≥32). (**J**) Blockage of VC synaptic transmission inhibits serotonin-induced egg laying in HSN-ablated animals. HSN-ablated *egl-1(986dm)* mutant animals expressing TeTx in the VCs were placed in M9 buffer or M9 containing 18.5 mM serotonin; asterisks indicate *p*≤0.0002, pound indicates *p*=0.0225 and n.s. indicates *p*>0.05 (Kruskal-Wallis test with Dunn’s correction for multiple comparisons; n≥22). (**K**) Animals with inhibited VC neurotransmission lay fewer eggs compared to wild-type animals in response to 18.5 mM serotonin under otherwise normal culturing conditions on solid media agar plates; asterisks indicate *p*<0.0001, pound indicates *p*=0.0026, and n.s. indicates *p*>0.05 (one-way ANOVA with Bonferroni’s correction for multiple comparisons; n≥10). (**L**) Measurement of steady-state egg accumulation in animals expressing TeTx in the VCs in an *nlp-3(tm3023)* mutant background (±95 confidence intervals for the mean); asterisks indicate *p*<0.0001 and n.s. indicates *p*>0.05 (one-way ANOVA with Bonferroni’s correction for multiple comparisons; n≥35). (**M**) Measurement of steady-state egg accumulation in animals expressing Histamine-gated chloride channels (HisCl) in either the VC or HSN neurons grown with or without histamine; asterisk indicates *p*<0.0001 and n.s. indicates *p*>0.05 (one-way ANOVA with Bonferroni’s correction for multiple comparisons; n≥33). (**N**) Acute electrical silencing of the VCs blocks serotonin-induced egg laying. Animals expressing HisCl in the VCs in either a wild-type or HSN-ablated *egl-1(986dm)* mutant background were incubated with 0 or 4 mM histamine for four hours, placed into wells with M9 buffer with 0 or 18.5 mM serotonin, and the number of eggs laid after 1 hour were counted; asterisks indicates *p*<0.05 and n.s. indicates *p*>0.05 (Kruskal-Wallis test with Dunn’s correction for multiple comparisons; n≥12 for control M9 and n≥24 for serotonin).

Here we address the function of the VC neurons during egg-laying behavior. We find that the VCs function within a serotonergic pathway to drive egg release. The VCs provide excitatory input to convert the initial stages of vulval muscle contraction into a successful egg-laying event. The VCs achieve this through direct activation in response to vulval muscle excitation and contraction, forming a positive feedback loop until successful egg laying is achieved.

## Results

The VCs are a group of six, hermaphrodite-specific, cholinergic motor neurons spaced out along the ventral cord that make synapses onto the HSNs, vm2 vulval muscles, body wall muscles, and various motor neurons involved in locomotion (Figure 1A; Cook et al., 2019; White et al., 1986). VC4 and VC5 in particular are the most proximal to the vulva and make extensive synapses onto the vm2 vulval muscles (Cook et al., 2019; White et al., 1986). Egg-laying events are always accompanied by a VC Ca^2+^ transient, but not all VC Ca^2+^ transients coincide with egg release (Collins et al., 2016). ACh released from VCs has been suggested to act through nAChRs including UNC-29 on the vulval muscles to drive contraction (Kim et al., 2001; Schafer, 2006), as well as through the muscarinic ACh receptor GAR-2 on the HSNs to inhibit egg laying (Bany et al., 2003; Fernandez et al., 2020). VC signaling has also been suggested to slow animal locomotion speed around egg-laying events (Collins et al., 2016), likely through the synapses it makes onto the body wall muscles and locomotion motor neurons (Cook et al., 2019; White et al., 1986). In all, VC signaling appears to be complex and function through multiple pathways to regulate egg-laying circuit activity and behavior.

The vulval muscles are comprised of four vm1-type and four vm2-type muscle cells which are all electrically coupled but receive distinct synaptic input (White et al., 1986). The vm2 vulval muscles are innervated by the HSN and VC neurons (Cook et al., 2019; White et al., 1986). The vm1s receive cholinergic input from single VA and VB motor neurons which are part the locomotion circuit (White et al., 1986; Zhen & Samuel, 2015). Egg laying occurs when all 8 vulval muscle cells are active and pull the vulva open to expel an egg (Zhang et al., 2008). Serotonergic input from the HSNs helps to excite the vm2 vulval muscles and coordinate their activity during egg laying (Brewer et al., 2019; Li et al., 2013; Shyn et al., 2003). The vm1 vulval muscles show rhythmic activity both during and outside of egg laying, suggesting that they are either intrinsically active or excited by VA and VB synaptic inputs (Collins et al., 2016; Collins & Koelle, 2013; Shyn et al., 2003). Vulval muscle Ca^2+^ transients start in the vm1 muscle cells and propagate into the vm2 muscles during egg laying events (Brewer et al., 2019; Collins & Koelle, 2013), suggesting vm1 excitation provides a trigger for full vulval muscle contraction and egg laying.

The pair of hermaphrodite-specific HSN neurons make synapses onto both the VCs and the vm2 vulval muscles (Cook et al., 2019; White et al., 1986). Serotonin and NLP-3 neuropeptides released by the HSNs have been shown to be critical for normal egg-laying behavior to occur (Brewer et al., 2019; Waggoner et al., 1998). Consistent with the role of HSNs initiating egg-laying behavior, exogenous serotonin potently increases the Ca^2+^ activity of the vulval muscles and VC neurons (Shyn et al., 2003; Zhang et al., 2008). Additionally, laser ablation experiments have indicated that loss of the VC neurons disrupts the induction of egg laying in response to serotonin (Shyn et al., 2003; Waggoner et al., 1998). Thus, the role of the VC neurons in the egg-laying circuit and behavior may be closely associated with serotoninergic signaling from the HSNs.

### The VC neurons promote egg laying in response to serotonin

To test directly how loss of synaptic transmission from the VC motor neurons affects egglaying circuit activity and behavior (Figure 1B), we used a modified *lin-11* promoter/enhancer (Bany et al., 2003) to drive transgenic expression of Tetanus Toxin (TeTx; Jose et al., 2007) which blocks both neurotransmitter and neuropeptide release (Whim et al., 1997) from the six VC neurons. Expression of TeTx in the VC neurons did not cause any gross defects in steady-state egg accumulation compared to non-transgenic control animals, indicating that the VC neurons are not strictly required for egg laying (compare Figures 1C and 1D, quantified in Figure1G). In contrast, *egl-1(n986dm)* mutant animals in which the HSNs undergo apoptosis (Trent et al., 1983), showed a dramatic impairment of egg laying, accumulating significantly more embryos (Figures 1E and 1G). Previous studies indicated that laser ablation of both the HSNs and VCs caused additive defects in egg laying (Waggoner et al., 1998). However, transgenic expression of TeTx in the VCs in HSN-deficient *egl-1(n986dm)* mutant animals did not significantly enhance their defects in egg laying (Figure 1F and 1G). We have previously shown that wild-type animals lay their first egg around 7 hours after the L4-adult molt, a time when the VCs show their first activity (Ravi et al., 2018a). Animals with inhibited VC neurotransmission showed no significant change in the onset of egg laying compared to non-transgenic control animals (Figure 1H). In contrast, the onset of egg laying in *egl-1(n968dm)* mutant animals lacking HSNs is significantly delayed, occurring about 18 hours after the L4-adult molt (Figure 1H). Expression of TeTx in the VCs in HSN-deficient animals did not enhance the delay in egg laying significantly (Figure 1H). These results together show that transmitter-mediated signaling from the VCs is not required for egg-laying behavior to occur under normal culturing conditions.

We next used a drug-treatment approach to explore possible functions of the VC neurons that may not be apparent in animals under standard laboratory conditions. Serotonin potently stimulates egg laying even in conditions where egg laying is normally inhibited, such as in liquid M9 buffer (Trent et al., 1983). As expected, serotonin promotes egg laying in M9 buffer in wild-type animals (Figure 1I). We found that transgenic animals expressing TeTx in the VCs failed to lay eggs in response to exogenous serotonin in M9 buffer across varying serotonin concentrations (Figure 1I). Going forward, we chose to conduct experiments at a single serotonin concentration of 18.5 mM. HSN-deficient *egl-1(n986dm)* mutant animals are egg-laying defective under normal conditions but will still lay eggs in response to exogenous serotonin (Schafer et al., 1996), and this response was suppressed in animals lacking VC neurotransmission at 18.5 mM serotonin (Figure 1J). This resistance to exogenous serotonin in VC neurotransmission-inhibited animals was not unique to M9 buffer, as transgenic animals placed on serotonin-infused agar also showed a reduced egg-laying response (Figure 1K).

These results show that VC neurotransmitter release serves an important role for egg laying in response to exogenous serotonin.

Despite the VCs being important for egg laying in response to exogenous serotonin (Figure 1I-K), animals with blocked VC neurotransmission are still able to lay eggs at a normal rate (Figure 1G). One potential explanation for this is that NLP-3 neuropeptide signaling is able to compensate for the loss of the VC-mediated serotonergic signaling pathway (Brewer et al., 2019). Based on this model, blocking VC neurotransmission in an *nlp-3* mutant background might phenocopy animals lacking all neurotransmitter release from the HSNs, as seen in the *egl-1(n986dm)* mutant. However, we found no significant increase in egg accumulation when blocking VC neurotransmission in an *nlp-3* mutant background (Figure 1L). This suggests that under standard *C. elegans* culture conditions, serotonin is able to signal through VC-independent pathway to retain a normal rate of egg laying, likely by acting on the vulval muscles directly (Hapiak et al., 2009).

It was possible that the observed defects in serotonin response caused by inhibiting VC neurotransmission could be due to impaired circuit development and/or by compensatory changes in circuit activity. Expression from the VC-specific promoter used to drive TeTx begins in the L4 stage as the egg-laying circuit is completing development, well before the onset of egg-laying behavior (Ravi et al., 2018a). To silence the VCs acutely after circuit development is complete, we transgenically expressed Histamine-gated chloride channels (HisCl) and treated the animals with exogenous histamine (Pokala et al., 2014; Ravi et al., 2018a). Histamine silencing of the VCs caused no gross changes in steady-state egg accumulation (Figure 1M), confirming the results with TeTx that VC neurotransmission is not required for egg laying. However, acute histamine silencing of the VCs reduced egg laying in response to serotonin in both wild-type and HSN-deficient *egl-1(n986dm)* mutant animals (Figure 1N), consistent with the results observed when blocking VC neurotransmission with TeTx. Together, these results show that both VC neuron activity and synaptic transmission are dispensable for egg laying under normal growth conditions, but the VCs do facilitate egg laying in response to exogenous serotonin.

### VC Ca^2+^ activity is coincident with vulval muscle activity and egg laying

The VC neurons show rhythmic Ca^2+^ activity during the egg-laying active state (Collins et al., 2016). However, only ∼1/3 of VC Ca^2+^ transients were temporally coincident with vulval muscle contractions that resulted in egg release (Collins et al., 2016), raising questions about the function of the VC Ca^2+^ transients that do not coincide with egg laying. To understand the function of VC Ca^2+^ activity, we expressed GCaMP5 in the VC neurons to measure Ca^2+^ dynamics in the VC cell bodies and processes most proximal to the vulva (VC4 & VC5; Figure2A and Figure 2B) while simultaneously observing vulval opening and egg-laying behavior in a separate brightfield channel, as described (Ravi et al., 2018b). Since *C. elegans* neurons generally exhibit graded potentials, measurements of VC Ca^2+^ near synapses can be expected to correspond closely with the degree of neurotransmitter and neuropeptide release (P. Liu et al., 2013; Q. Liu et al., 2009). As expected, we found that egg-laying events were always accompanied by a VC Ca^2+^ transient, but not every VC Ca^2+^ transient resulted in an egg-laying event (Figure 2C; Movie 1). These remaining VC Ca^2+^ transients were almost always observed with weak muscle contraction and partial opening of the vulva, termed a vulval muscle “twitch,” which are primarily mediated by the vm1 vulval muscles (Figure 2C and 2F; Collins & Koelle, 2013). As quantified in Figure 2F, we found that 48% of VC Ca^2+^ transients were associated with vulval muscle twitch contractions, and 39% of VC Ca^2+^ transients were associated with strong vulval muscle contractions that support egg release, which are the result of simultaneous vm1 and vm2 vulval muscle contraction (Collins & Koelle, 2013). To determine if presynaptic HSN activity was sufficient to drive VC and vulval muscle activity downstream, we optogenetically stimulated HSNs transgenically expressing Channelrhodopsin-2 and simultaneously recorded VC Ca^2+^ activity and vulval muscle contractility (Figure 2A). As previously reported, optogenetic HSN stimulation was sufficient to induce an active egg-laying behavior state (Collins et al., 2016). We found that optogenetic stimulation of HSNs drove a robust increase in VC Ca^2+^ transients that were associated with egg-laying events, but we still observed VC Ca^2+^ transients during the weaker vulval muscle twitching contractions (Figure 2D). HSN optogenetic stimulation rapidly elevated average Ca^2+^ levels in the VCs, within 5 seconds after blue light exposure (Figure 2E).

**Figure 2.**
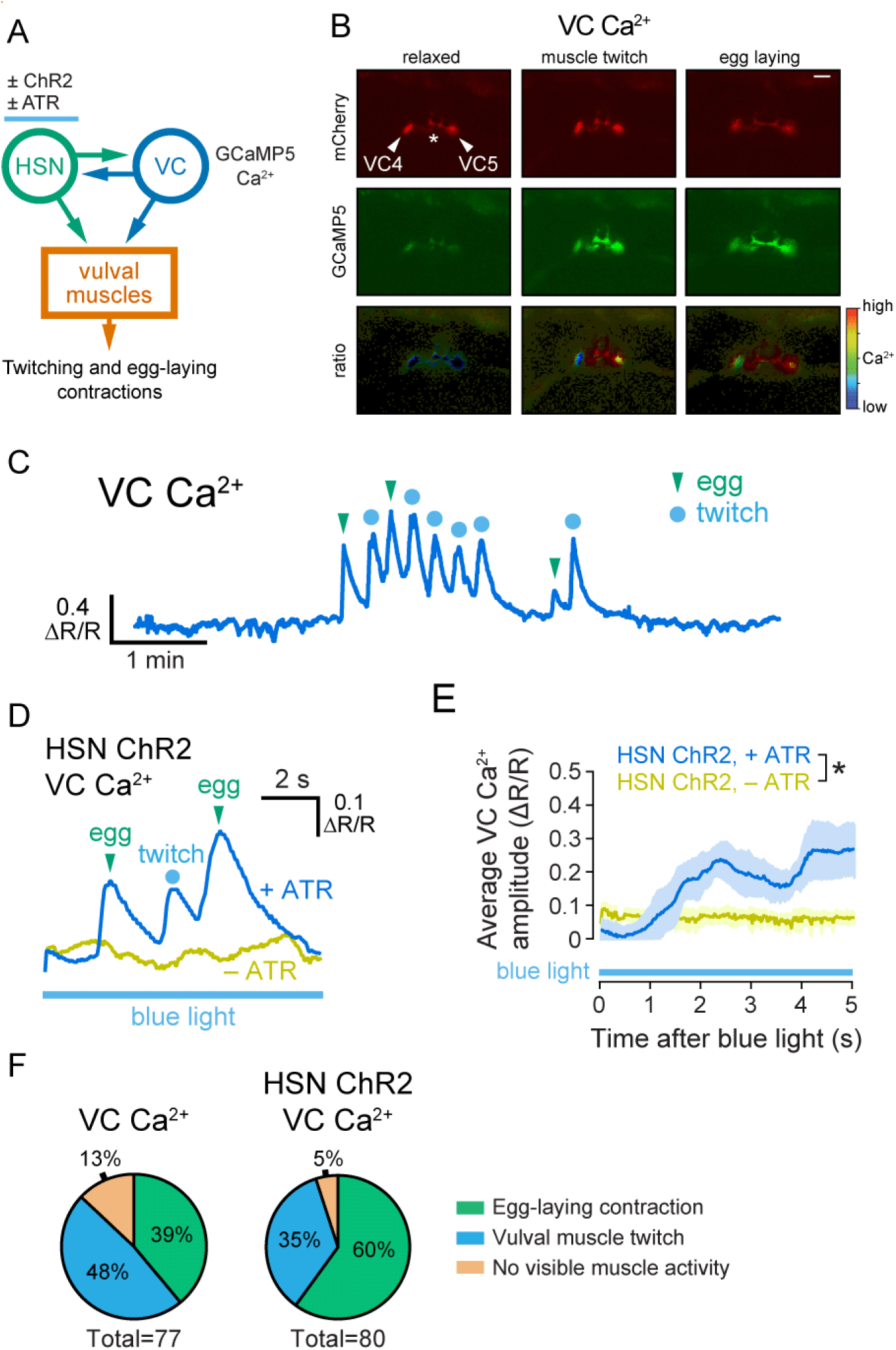
The VC neurons are active during both weak vulval muscle twitching and strong egg-laying contractions. (**A**) Cartoon of the circuit and experimental manipulations. GCaMP5 was expressed in the VC neurons to record Ca^2+^ activity, and Channelrhdopsin-2 was expressed in HSNs to provide optogenetic stimulation of egg laying. (**B**) Representative still images of VC mCherry, GCaMP5, and GCaMP5/mCherry ratio (ΔR/R) during inactive and active egg-laying behavior states. Asterisk indicates vulva. Scale bar is 20 μm. (**C**) Representative trace of VC GCaMP5/mCherry Ca^2+^ ratio in freely behaving, wild-type animals during an egg-laying active state. (**D**) Representative trace of VC GCaMP5/mCherry Ca^2+^ ratio after optogenetic stimulation of ChR2 in the HSN neurons in animals grown in the absence (-ATR, top) and presence (+ATR, bottom) during 10 s of continuous blue light exposure. (**E**) Average vulval muscle Ca^2+^ levels (mean ±95% confidence intervals; asterisk indicates *p*<0.0001, Student’s t test; n=10). (**F**) Graph showing vulval muscle contractile activity for each VC Ca^2+^ transient during endogenous egg-laying active states (left) or in response to HSN optogenetic stimulation (right).

The light-dependent increase in VC Ca^2+^ activity and vulval muscle contractions were not observed in animals grown without the essential cofactor, all-trans-retinal (ATR; Figure 2D and 2E). During optogenetic HSN activation, 60% of VC Ca^2+^ transients were associated with egg-laying events while only 35% were associated with vulval muscle twitches, a significant difference from control animals not subjected to HSN optogenetic stimulation that we attribute to an increase in egg laying frequency (Figure 2F). Because VC Ca^2+^ activity rises at the same time as vulval opening and prior to egg release, these results are consistent with either a model where the VCs promote vulval muscle contractility, or a model where the VCs are activated in response to downstream vulval muscle contraction.

### The VC neurons promote vulval muscle activity and contraction

The VC motor neurons synapse onto the vulval and body wall muscles where they are thought to release ACh to drive contraction (Duerr et al., 2008; White et al., 1986; Zhang et al., 2008). While VC activity and neurotransmission is not required for egg laying (Figure 1G and1K; Laura E. Waggoner et al., 1998), the VCs may still release ACh to regulate vulval muscle contractility. To test if the VCs can excite the vulval muscles directly, we expressed Channelrhodopsin-2 in the VC neurons and performed ratiometric Ca^2+^ imaging in the vulval muscles after exposure to blue light (Figure 3A; Movie 2). Optogenetic stimulation of the VCs led to an acute induction of vulval muscle Ca^2+^ activity within 5 s but was unable to drive full vulval muscle contractions and egg release (Figure 3B and 3C). However, average vulval muscle Ca^2+^ transient amplitude after optogenetic stimulation was not significantly higher through the duration of the recording (Figure 3D). Vulval muscle Ca^2+^ transient frequency was reduced, which may result from the increased duration of each individual transient (Figures 3E and 3F). This result shows that VC activity alone is not able to maximally excite the vulval muscles to the point of egg laying, but that VC activity is excitatory and can sustain ongoing vulval muscle Ca^2+^ activity. We find these results consistent with a model where serotonin and NLP-3 neuropeptides released from the HSNs signal to enhance the excitability and contractility of the vulval muscles for egg laying (Brewer et al., 2019), while the ACh released from the VCs prolong the contractile phase to facilitate egg release.

**Figure 3.**
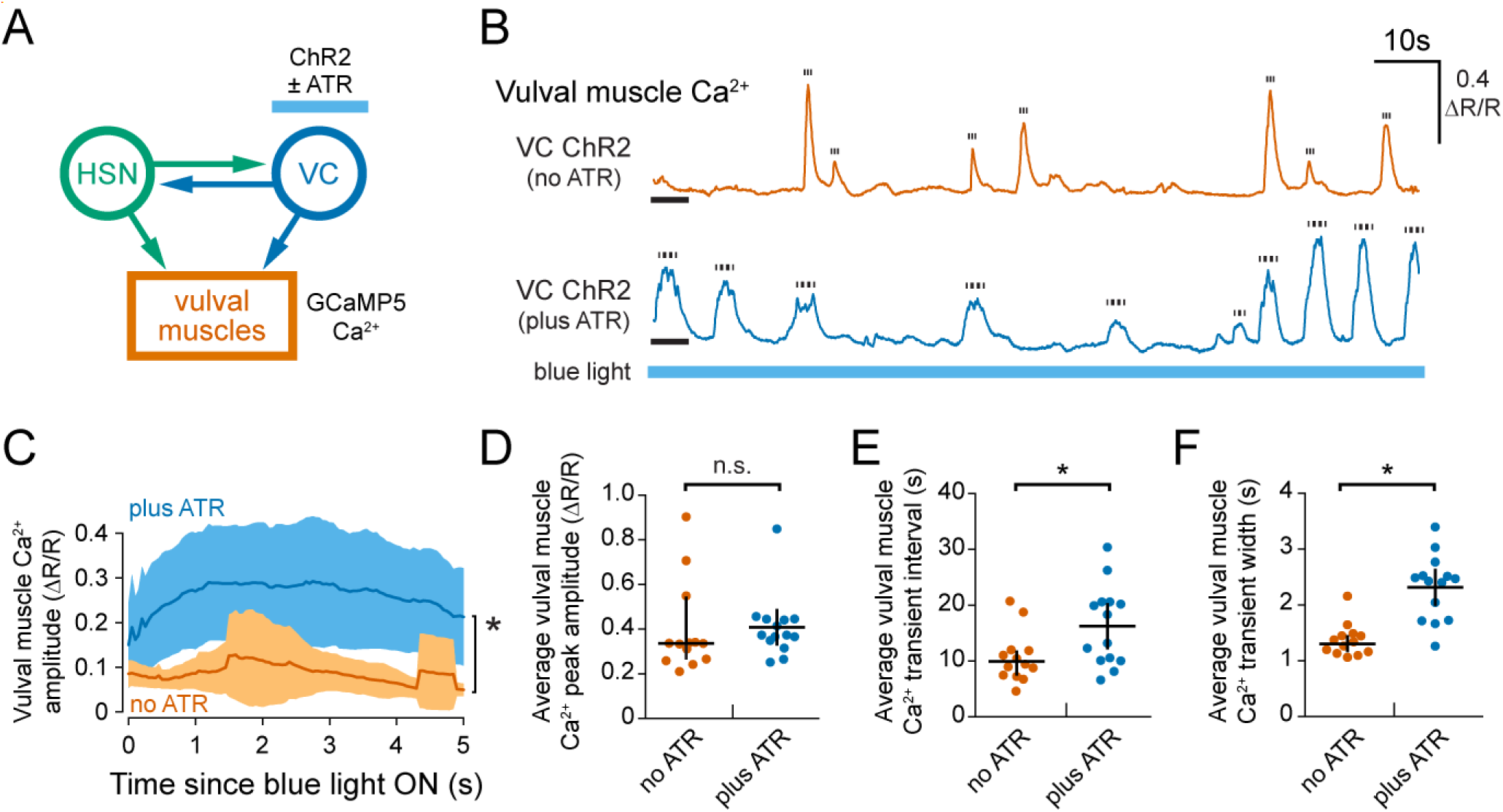
Optogenetic VC activation induces and sustains vulval muscle Ca^2+^ activity. **(A)** Cartoon of circuit and experiment. Channelrhdopsin-2 was expressed in the VC neurons and GCaMP5 was expressed in the vulval muscles. (**B**) Representative traces of vulval muscle GCaMP5/mCherry Ca^2+^ ratio (ΔR/R) in animals grown in the absence (-ATR, top) or presence (+ATR, bottom) in response to 5 minutes of continuous blue light exposure. Tick marks show duration at half-maximal amplitude of each measured Ca^2+^ transient. Black bar indicates first 5 seconds following blue light exposure. (**C**) Averaged vulval muscle Ca^2+^ transient activity (shaded region indicates ±95% confidence intervals) during the first 5 seconds of optogenetic activation of the VC neurons in -ATR (control; blue) and +ATR conditions (orange; asterisk indicates *p*=0.003, Student’s t test; n≥13 animals). (**D**) Scatterplot showing the average peak amplitude of vulval muscle Ca^2+^ transients (±95% confidence intervals) per animal in response to VC optogenetic stimulation during 5 minutes of continuous blue light (n.s. indicates *p*>0.05; Student’s t test; n≥13 animals). (**E**) Scatterplot showing the average time between vulval muscle Ca^2+^ transients (±95% confidence intervals) per animal during VC optogenetic stimulation during 5 minutes of continuous blue light (asterisk indicates *p*=0.0236, Student’s t test; n≥13 animals). (**F**) Scatterplot showing the average vulval muscle Ca^2+^ transient width (±95% confidence intervals) per animal during VC optogenetic stimulation during 5 minutes of continuous blue light (asterisk indicates *p*<0.0001, Student’s t test; n≥13 animals; error bars indicate 95% confidence intervals for the mean).

### The VCs facilitate successful vulval opening during egg laying

Loss of VC activity or synaptic transmission caused a specific defect in serotonin-induced egg laying, suggesting the VCs are required for proper vulval muscle Ca^2+^ activity and/or contractility. We recorded vulval muscle Ca^2+^ activity in freely behaving animals transgenically expressing TeTx in the VCs (Figure 4A; Movie 3). Vulval muscle Ca^2+^ activity in wild-type animals is characterized by low activity during the ∼20 minute egg-laying inactive state, and periods of high activity during the ∼2 minute egg-laying active state (Figure 4B; Collins et al., 2016; Laura E. Waggoner et al., 1998). Expression of TeTx in the VCs did not significantly affect the overall frequency or amplitude of vulval muscle Ca^2+^ transients during egg-laying inactive states compared to wild-type control animals (Figures 4C and 4D). However, inhibition of VC neurotransmission led to larger amplitude Ca^2+^ transients during the active phase (Figure 4D). Since previous work suggested the VCs release ACh that inhibits egg laying (Bany et al., 2003), the simplest explanation for this phenotype would be that the VCs normally inhibit vulval muscle Ca^2+^ activity. However, our results show optogenetic activation of the VCs increased vulval muscle Ca^2+^ transient duration with minimal effect on amplitude (Figures 3C, 3D, and 3F). Further inspection of vulval muscle Ca^2+^ traces of individual animals showed that transgenically expressing TeTx in the VCs had large vulval muscle Ca^2+^ transients of amplitude similar to egg-laying Ca^2+^ transients (>1.0 ΔR/R), but without an egg being successfully released (Figure 4Band 4E; Movie 3). These large amplitude transients that did not lead to egg laying were thus termed “failed egg-laying events.” Such failed egg-laying events were infrequent in vulval muscle Ca^2+^ recordings from wild-type animals, but they occurred more frequently than successful egg-laying events in transgenic animals expressing TeTx in the VCs (Figure 4E). Based on these results, it appears that VC neurotransmission does not initiate vulval muscle Ca^2+^ transients but is instead critical for coordinating vulval muscle Ca^2+^ activity and contraction across the vulval muscle cells to allow for successful egg release.

**Figure 4.**
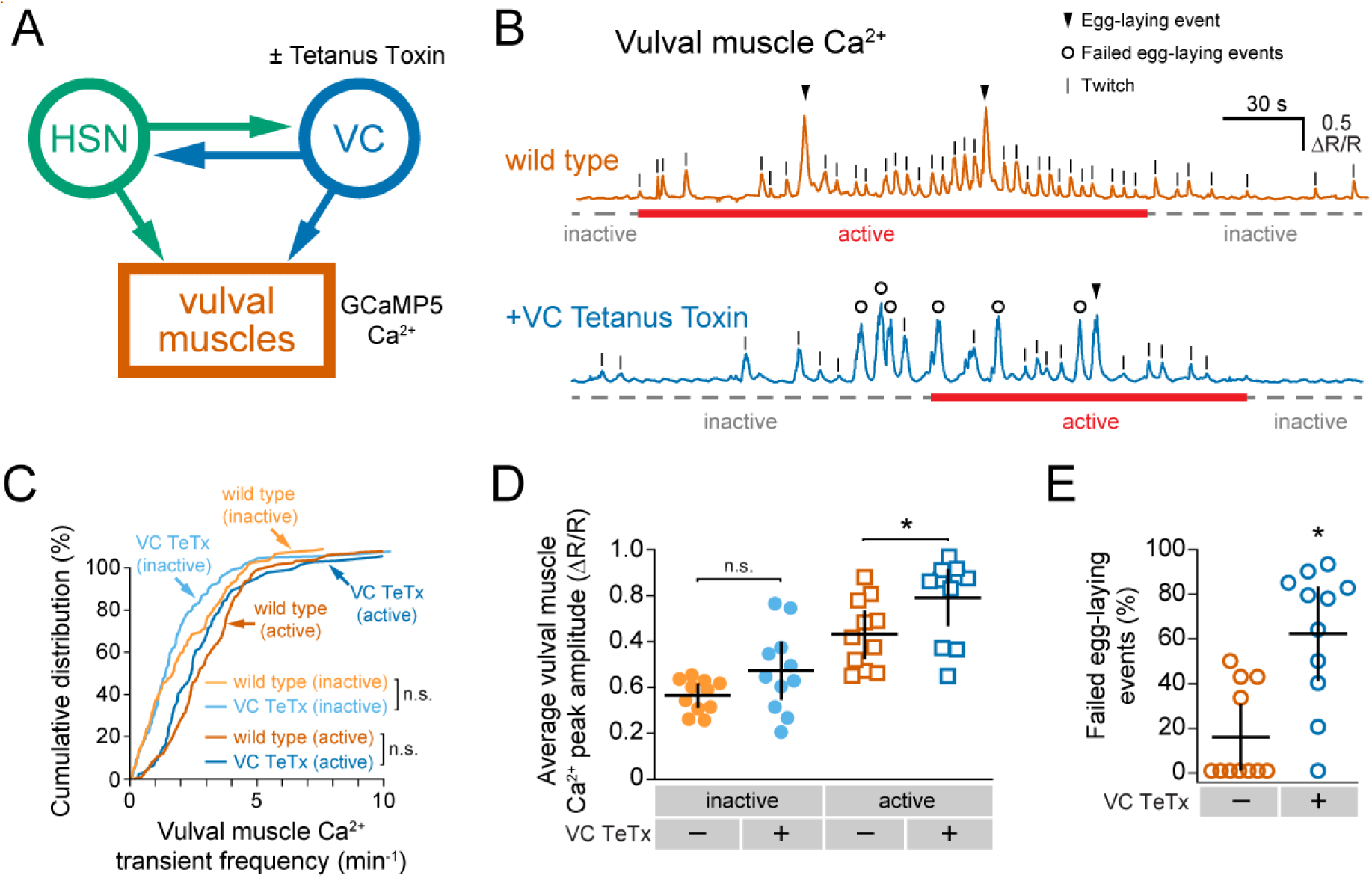
Blocking VC neurotransmission reduces the success rate of egg laying. **(A)** Cartoon of circuit and experiment. TeTx was expressed in the VC neurons to block their neurotransmitter release and GCaMP5 was expressed in the vulval muscles to record Ca^2+^ activity. (**B**) Representative traces of vulval muscle GCaMP5/mCherry Ca^2+^ ratio (ΔR/R) in wild-type (top) and TeTx expressing transgenic animals (bottom). Arrowheads indicate successful egg-laying events, and open circles indicate strong (>1.0 ΔR/R) vulval muscle Ca^2+^ transients of “failed egg-laying events.” Egg-laying behavior state (inactive or active) duration is indicated below each trace. (**C**) Cumulative distribution plots of all vulval muscle Ca^2+^ transients across wild-type and transgenic animals expressing TeTx in the VCs. Transients were parsed into egg-laying active and inactive phases (n.s. indicates p>0.05, Kruskal-Wallis test with Dunn’s correction for multiple comparisons; n≥200 transients from 11 animals). (**D**) Average vulval muscle Ca^2+^ transient peak amplitudes per animal during the inactive and active egg-laying phase (asterisk indicates *p*=0.0336, one-way ANOVA with Bonferroni’s correction for multiple comparisons; n=11 animals). (**E**) Percentage of failed egg-laying events in wild-type and transgenic animals expressing TeTx in the VCs (asterisk indicates *p*=0.0014, Mann-Whitney test; n=11 animals).

Egg laying occurs when strong vulval muscle Ca^2+^ activity drives the synchronous contraction of all the vulval muscle cells (Brewer et al., 2019; Li et al., 2013) that allows for the mechanical opening of the vulva in phase with locomotion for efficient egg release (Collins et al., 2016; Collins & Koelle, 2013). Egg-laying events are characterized by coordinated Ca^2+^ activity between the vm1 vulval muscles that extend to the ventral tips of the vulva and the medial vm2 vulval muscles, leading to full contraction and egg release (Figure 5A). This Ca^2+^ activity is distinct from weak vulval muscle twitching contractions that are confined to the vm1 muscles (Figure 5A; Collins & Koelle, 2013). Failed egg-laying events exhibit Ca^2+^ activity more similar to egg-laying events, where Ca^2+^ signal is high across both vm1 and vm2 muscles (Figure 5A). To understand why egg laying was less likely to occur during strong vulval muscle Ca^2+^ transients in VC neurotransmission-inhibited animals, we measured features of contractile events during egg laying. Contraction can be directly quantified by measuring the reduction in muscle area in fluorescent micrographs, and vulval opening can be quantified by measuring the changing distance between the vulval muscle cells positioned anterior and those posterior to the vulval slit (Figure 5A). During egg-laying events in wild-type animals, a strong cytosolic Ca^2+^ transient correlates with a ∼50 µm^2^ contraction of muscle size followed by a rebound phase after egg release (Figure 5B, top and middle). Simultaneously, the anterior and posterior muscles separate by ∼10 µm, facilitating egg release (Figure 5B, bottom). We found differences in the kinetics and extent of vulval muscle opening between wild-type and TeTx-expressing transgenic animals during successful egg-laying events (compare Figures 5B and 5C). The vulval muscles opened wider and the degree of contraction was greater during egg-laying events in animals where VC neurotransmission was inhibited with TeTx (Figure 5C), possibly because the vulval muscle Ca^2+^ rise started earlier in these animals. During failed egg-laying events, the vulval muscles showed only modest contraction and only separated by ∼5 µm, insufficient to allow egg release, despite reaching similar levels of cytosolic Ca^2+^ (Figure 5D). Animals with blocked VC neurotransmission exhibited even less separation of the vulval muscles during failed egg-laying events (compare Figures 5D and 5E). In contrast to wild type animals, animals with inhibited VC neurotransmission have many more failed egg-laying events during which they exhibit vulval opening kinetics more similar to twitches (compare Figure 5F and 5G). To understand the relationship between vulval muscle Ca^2+^ levels and vulval opening, we measured the distance between anterior and posterior vulval muscles during weak twitching, failed egg-laying events, and successful egg-laying events (Figure 5F). In both wild-type and TeTx-expressing animals, we noted a linear relationship of low but positive slope between vulval opening and Ca^2+^ levels <1.0 ΔR/R (Δopening / ΔCa^2+^) during weak twitching contractions (Figure 5F and 5G). However, as Ca^2+^ levels rose above 1.0 ΔR/R, the muscles reached threshold for full opening, allowing successful egg release (Figure 5F). The steep, linear Δopening / ΔCa^2+^ relationship during failed egg-laying events suggests a threshold of Ca^2+^ drives an all-or-none transition to full contraction, vulval opening, and egg release. In VC neurotransmission-inhibited animals, the shallow Δopening / ΔCa^2+^ relationship continued as weak twitches transitioned into failed egg-laying events, with many strong vulval muscle Ca^2+^ transients failing to open the vulva sufficiently for egg release (Figure 5G). However, successful egg-laying events in VC neurotransmission-inhibited animals still showed a sharp threshold between Ca^2+^ levels and the degree of vulval opening. This raises the possibility of two types of failed egg-laying events: one that is shared with wild-type animals, and another that is unique to animals with inhibited VC neurotransmission.

**Figure 5.**
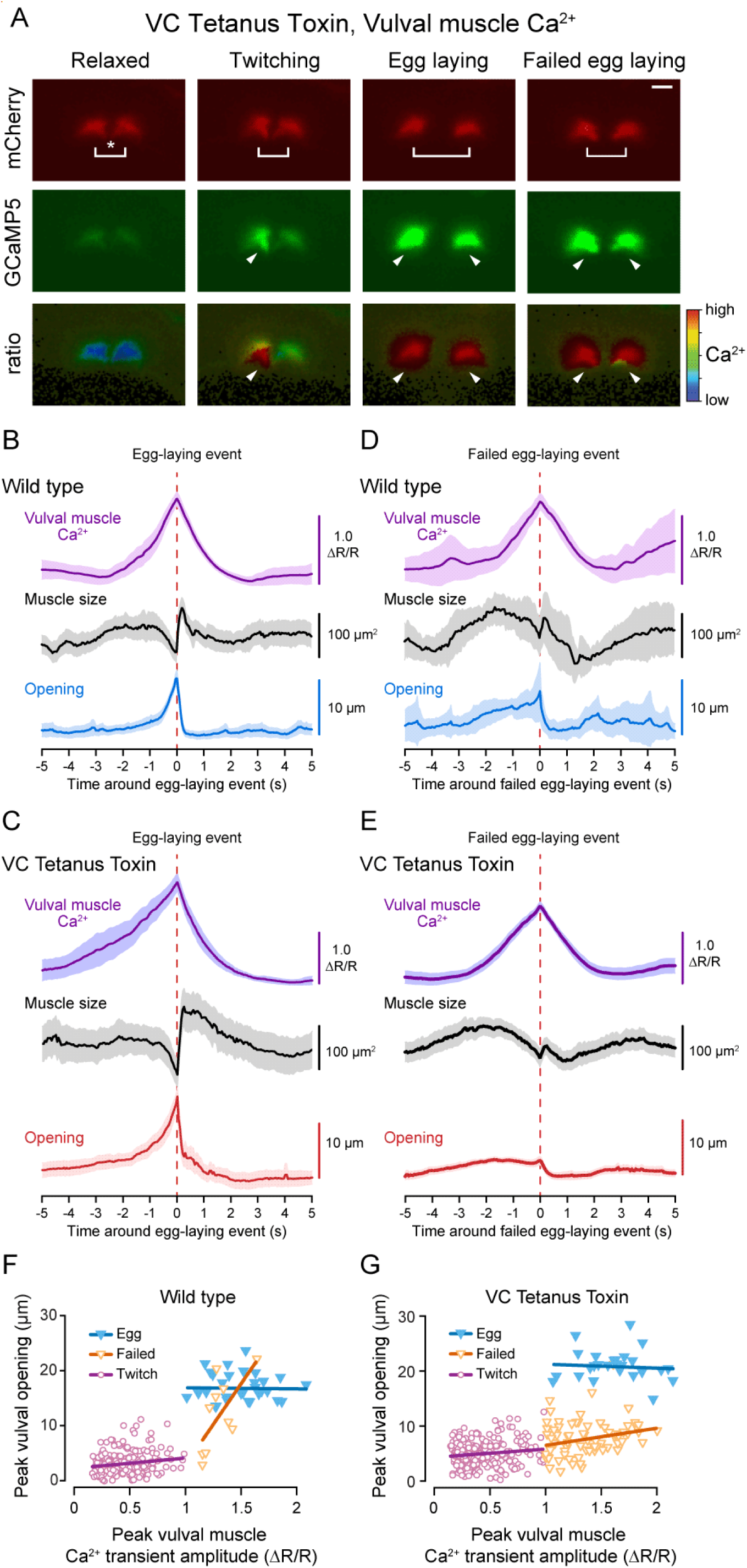
Blocking VC neurotransmission decouples vulval muscle Ca^2+^activity and egg laying. **(A)** Vulval muscle sizes and distances were quantified by measuring the changes in area and centroid of the anterior and posterior muscle groups in mCherry channel micrographs during GCaMP5/mCherry ratiometric imaging. Shown are still images of the vulval muscles during representative muscle states of relaxation, twitching, egg laying, and failed egg-laying events. Brackets indicate the distance between the centroid of vulval muscle halves, arrowheads indicate high Ca^2+^ activity, and asterisk indicates vulva. Scale bar is 20 μm. (**B-D)** Mean traces (±95 confidence intervals) of vulval muscle Ca^2+^ (GCaMP5/mCherry ratio), vulval muscle area (µm^2^), and vulval muscle centroid distance (µm) during successful (B and C; n≥26 from 11 animals) and failed egg-laying events (D and E; n≥10 from 11 animals) in wild-type (B and D) and transgenic animals expressing TeTx in the VC neurons (C and E). (**E**-**F**) Scatter plot showing peak vulval muscle Ca^2+^ amplitude in relation to the corresponding vulval muscle opening distance in wild-type (E) and transgenic animals expressing TeTx in the VC neurons (F). Lines through points represent simple linear regression for each labeled grouping.

### VC neurotransmission regulates HSN command neuron and egg-laying circuit activity

To determine whether VC synaptic transmission regulates egg laying via HSN, we recorded HSN Ca^2+^ activity in wild-type and transgenic animals expressing TeTx in the VCs (Figure 6A). During the egg-laying active state, the HSNs drive egg laying during periods of increased Ca^2+^ transient frequency in the form of burst firing (Figure 6B; Collins et al., 2016; Ravi et al., 2018a). We observed a significant increase in HSN Ca^2+^ transient frequency when VC synaptic transmission was blocked compared to non-transgenic control animals (Figure 6C). Wild-type animals spent ∼11% of their time exhibiting high frequency burst activity in the HSN neurons, while transgenic animals expressing TeTx in the VC neurons spent ∼21% of their time exhibiting HSN burst firing activity (Figure 6D). These results are consistent with the interpretation that VC neurotransmission is inhibitory toward the HSNs such as proposed in previous studies (Bany et al., 2003; Zhang et al., 2008), but the steady-state egg accumulation of animals expressing VC-specific TeTx or HisCl is normal (Figure 1G, K). We have previously shown that HSN burst firing is regulated by egg accumulation and feedback of vulval muscle Ca^2+^ activity (Ravi et al., 2018a), which could be enhanced by the high rate of failed egg-laying events observed in VC TeTx transgenic animals (Figure 5E). We propose that the increased HSN burst firing seen in VC TeTx transgenic animals does not solely result from loss of inhibitory VC input, but also reflects the circuit prolonging the active state to compensate for more failed egg-laying transients in the absence of excitatory VC input.

**Figure 6.**
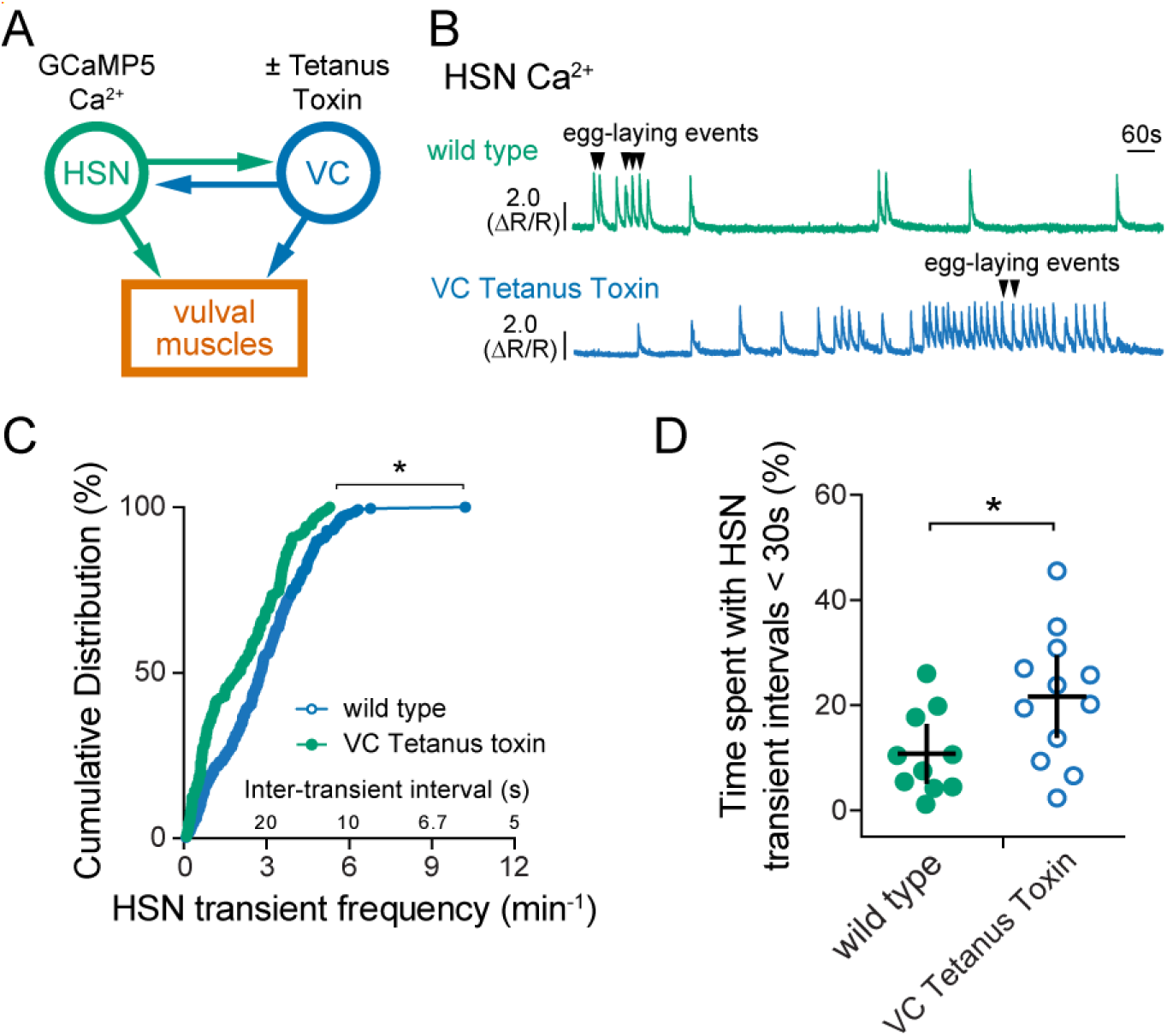
Blocking VC neurotransmission increases HSN Ca^2+^ activity. **(A)** Cartoon of circuit and the experiment. TeTx was expressed in the VC neurons to block their neurotransmitter release and HSN Ca^2+^ activity was recorded through GCaMP5 imaging. (**B**) Representative traces of HSN neuron GCaMP5/mCherry Ca^2+^ ratio (ΔR/R) in a wild-type background (top, green) and animals expressing TeTx in the VCs (bottom, blue). (**C**) Cumulative distribution plots of the instantaneous frequency of HSN Ca^2+^ transients in wild-type and TeTx-expressing animals (asterisk indicates p=0.0006, Kolmogorov-Smirnov test; n≥154 from ≥10 animals). (**D**) Scatter plot for the percentage of time each animal spent with HSN Ca^2+^ inter-transient intervals that were less than 30 seconds, an indicator of hyperactivity (asterisk indicates *p*=0.00272, Student’s test; n≥10; error bars indicate 95% confidence intervals for the mean).

### The VC motor neurons are responsive to vulval muscle activity and contraction

VC Ca^2+^ activity is coincident with strong vulval muscle twitching and egg-laying contractions (Figure 2; Collins et al., 2016). In addition to making synapses onto the vm2 muscles whose contraction drives egg laying, the VCs extend neurites along the vulval hypodermis devoid of synapses (White et al., 1986), suggesting the VCs may respond to vulval opening. To test this model, we sought to induce vulval opening independent of endogenous circuit activity and presynaptic input from the HSNs. We transgenically expressed Channelrhodopsin-2 specifically in the vulval muscles using the *ceh-24* promoter to stimulate the vulval muscles and used GCaMP5 to record blue-light induced changes in vulval muscle Ca^2+^ activity (Figure 7A). Optogenetic stimulation of the vulval muscles triggered an immediate rise in vulval muscle cytosolic Ca^2+^, tonic contraction of the vulval muscles, vulval opening, and egg release (Figures 7B and 7C). Even though optogenetic stimulation resulted in sustained vulval muscle Ca^2+^ activity and contraction, vulval opening and egg release remained rhythmic and phased with locomotion, as previously observed in wild-type animals (Collins et al., 2016; Collins & Koelle, 2013). Simultaneous bright field recordings showed the vulva only opened for egg release when the adjacent ventral body wall muscles were in a relaxed phase (Movie 4). We have previously shown that eggs are preferentially released when the vulva is at a particular phase of the body bend, typically as the ventral body wall muscles anterior to the vulva go into a more relaxed state (Collins et al., 2016; Collins & Koelle, 2013). We now interpret this phasing of egg release with locomotion as evidence that vulval muscle Ca^2+^ activity drives contraction, but the vulva only opens for successful egg release when contraction is initiated during relaxation of the adjacent body wall muscles. Together, these results show that optogenetic stimulation of the vulval muscles is sufficient to induce vulval muscle Ca^2+^ activity for egg-release in a locomotion phase-dependent manner.

**Figure 7.**
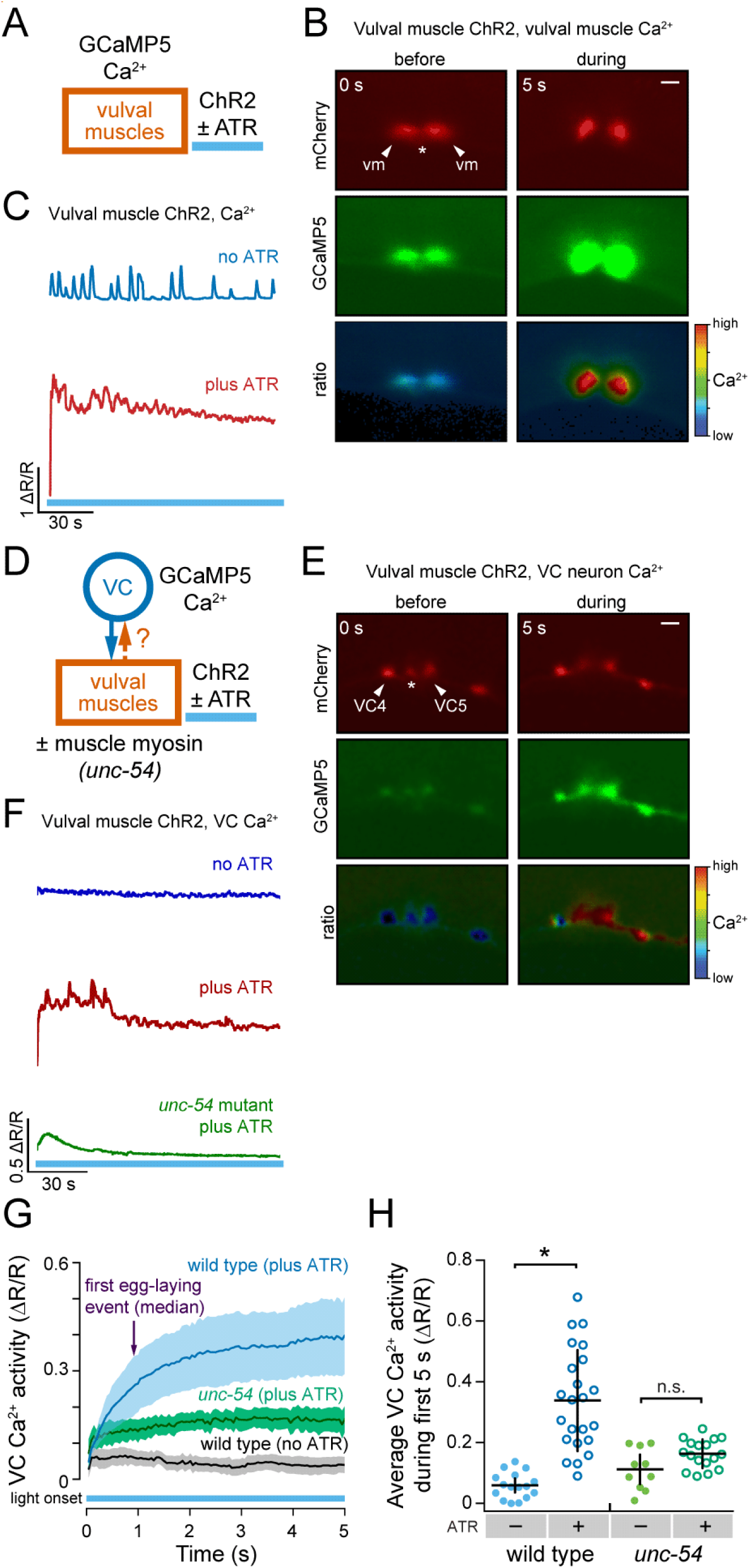
Optogenetic activation and contraction of the vulval muscles drives VC neuron activity. **(A)** Cartoon of experiment. Channelrhodopsin-2 and GCaMP5 were expressed in the vulval muscles to monitor cytosolic Ca^2+^ after optogenetic stimulation. (**B**) Representative still images of vulval muscle mCherry, GCaMP5, and GCaMP5/mCherry ratio during optogenetic activation of the vulval muscles. Arrowheads indicate each vulval muscle half, and asterisk indicates vulva. Scale bar is 20 μm. (**C**) Representative traces of vulval muscle GCaMP5/mCherry Ca^2+^ ratio (ΔR/R) in animals expressing ChR2 in the vulval muscles in the absence (-ATR, top) or presence (+ATR, bottom) of all-*trans* retinal during 3 minutes of continuous blue light exposure. (**D**) Cartoon of circuit and experiment. Channelrhdopsin-2 was expressed in the vulval muscles and GCaMP5 was expressed in the VC neurons to record Ca^2+^ activity in wild-type or *unc-54* myosin null mutant animals. (**E)** Representative still images of VC neuron mCherry, GCaMP5, and GCaMP5/mCherry ratio during optogenetic activation of the vulval muscles. Arrowheads indicate VC neuron cell bodies, and asterisk indicates vulva. Scale bar is 20 μm. (**F**) Representative traces of VC neuron GCaMP5/mCherry Ca^2+^ ratio (ΔR/R) in animals expressing ChR2 in the vulval muscles in the absence (-ATR, top) or presence (+ATR, bottom) of all-*trans* retinal during 3 minutes of continuous blue light exposure. **(G)** Averaged VC Ca^2+^ responses during the first 5 s. Error bands represent ±95 confidence intervals for the mean; n≥10 animals. (**H)** Scatter plot showing the average VC Ca^2+^ response during the first 5 seconds of optogenetic stimulation of the vulval muscles. Asterisk indicates *p*<0.0001; n.s. indicates p>0.05 (one-way ANOVA with Bonferroni’s correction for multiple comparisons; n≥10 animals).

To test the hypothesis that the VCs respond to vulval muscle activation, we recorded changes in VC Ca^2+^ upon optogenetic stimulation of the vulval muscles (Figure 7D). We observed a robust induction of VC Ca^2+^ activity upon blue light illumination (Figures 7E and 7F). The rise in VC Ca^2+^ occurred before the first egg-laying event, suggesting that this process is dependent on muscle activity and not necessarily the passage of an egg (Figure 7G; Movie 5).

This result demonstrates that the VCs can become excited in response to activity of the postsynaptic vulval muscles.

How do the vulval muscles activate the VCs? The VCs make both chemical and electrical synapses onto the vm2 vulval muscles (Cook et al., 2019; White et al., 1986). Depolarization of the vm2 vulval muscles might be expected to electrically propagate to the VC and trigger an increase in VC Ca^2+^ activity. Another possibility is that the VCs are mechanically activated in response to vulval muscle contraction and/or vulval opening. To test these alternate models, we optogenetically stimulated the vulval muscles of *unc-54* muscle myosin mutants, which are unable to contract their muscles, and recorded VC Ca^2+^ activity (Figure 7D). We found optogenetic activation of the vulval muscles failed to induce VC Ca^2+^ activity in *unc-54* mutants compared to the wild-type background (Figures 7G and 7H). While *unc-54* mutants appear to show some increase in VC Ca^2+^ activity following blue light stimulation of the vulval muscles, this increase was not statistically significant, suggesting indirect excitation of the VCs through gap junctions is insufficient on its own to induce robust VC Ca^2+^ activity. Together, these results support a model where the VC neurons are mechanically activated in response to vulval muscle contraction. Mechanical activation appears to drive VC activity and is mediated through the VC4 and VC5 neurites which are most proximal to the vulval canal through which eggs are laid.

## Discussion

The connectome of *C. elegans* has greatly informed neural circuit studies and contributed to studies revealing that connectivity alone is not sufficient to explain nervous system operations (Bargmann, 2012; Bentley et al., 2016). In the present study, we examined the neural circuit driving egg-laying behavior in *C. elegans* at a cellular resolution to reveal functional pathways and elements of the behavior that had not been discernable through connectome or prior genetic studies (Figure 8). We show that the cholinergic VC motor neurons contribute to egg laying in a serotonergic pathway. Our data show that HSN activity acts to excite the vulval muscles and VCs, and that VC Ca^2+^ activity is closely associated with visible vulval muscle contraction. In the absence of HSN-mediated potentiation, the VCs are able to excite the vulval muscles, but not to the threshold required for egg laying. This sub-threshold interaction is consistent with a model where serotonin-mediated potentiation of the VCs and vulval muscles is required before the VCs can facilitate egg laying through timely excitatory input. In the absence of VC neurotransmission, the HSNs and the vulval muscles show excess Ca^2+^ activity, but this excess activity does not correspond to increased egg laying. Instead, we find that the vulval muscles are less efficient at opening the vulva, indicating the VCs have a role in facilitating vulval muscle contraction. Surprisingly, optogenetic activation and contraction of the postsynaptic vulval muscles is sufficient to drive presynaptic VC Ca^2+^ activity. We propose that serotonin released from the HSNs signals to promote both vulval muscle contractility and VC sensitivity to that contraction. Following this potentiation, vulval muscle twitch contractions are able to mechanically activate the VCs to release excitatory ACh as part of a positive-feedback loop until the vulval muscles are fully contracted and egg laying occurs.

**Figure 8.**
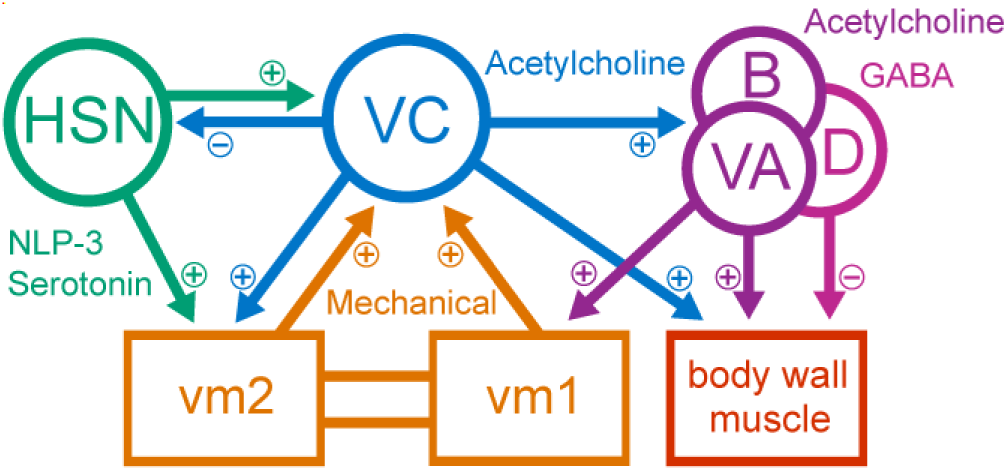
The VC neurons function to coordinate synchronized contraction of the vulval muscles for egg laying in response to serotonin-mediated potentiation. Working model of the functional connectivity within the *C. elegans* egg-laying circuit (Cook et al., 2019; White et al., 1986). Plus and minus signs indicate excitatory and inhibitory input, respectively. HSN command neurons release serotonin and NLP-3 to potentiate the VCs and vm2 vulval muscles (Figure 1 and Figure 2; Collins et al., 2016; Shyn et al., 2003; Waggoner et al., 1998; Zhang et al., 2008). VC neurons release acetylcholine that excites the vm2 vulval muscles, body wall muscles, and the VA/VB/VD locomotion motor neurons (Figure 3; Collins et al., 2016; Kim et al., 2001), as well as acetylcholine to inhibit the HSNs (Figure 6; Bany et al., 2003; Zhang et al., 2008). The VA/VB/VD neurons release acetylcholine and GABA onto the body wall muscles to regulate locomotion (Richmond & Jorgensen, 1999; Zhen & Samuel, 2015), and onto the vm1 vulval muscles to initiate contractions for egg laying (Collins et al., 2016; Collins & Koelle, 2013). Contraction of the vm1 vulval muscles electrically excites the vm2 vulval muscles and mechanically activates the VCs (Figure 7), forming a positive feedback loop until all vulval muscle cells are contracted and egg laying occurs.

Many behaviors require downstream feedback to help fine-tune movements and make adjustments based on the changing internal and external environment, such as with wing-beat patterns in *Drosophila* or head-eye coordination in humans (Bartussek & Lehmann, 2016; Fang et al., 2015). We find that the downstream target of the egg-laying neural circuit, the vulval muscles, signals upstream to facilitate proper completion of the behavior. We postulate that such feedback signaling is critical for two reasons. First, it can act to feedback inhibit the circuit to signal when the behavior has been executed. We find that vulval muscle contraction activates the VCs (Figure 7), which could contribute to inhibition of the egg-laying circuit through release of ACh acting on metabotropic receptors such as GAR-2 on the HSNs and vulval muscles (Bany et al., 2003; Fernandez et al., 2020; Zhang et al., 2008, 2010). GAR-2 has previously been shown to act in parallel with ionotropic receptors to differentially modulate locomotion behavior, providing both short- and longer-term effects in response to cholinergic signaling (Dittman & Kaplan, 2008; Zhen & Samuel, 2015). Second, feedback signaling could create a positive feedback loop to facilitate full execution of a behavior. In this circuit model, the VCs are mechanically activated by vulval muscle twitches which in turn leads to the VCs further exciting the vulval muscles until ubiquitous and coordinated contraction is achieved. A similar type of feedback activity has been demonstrated in the *C. elegans* SMDD neurons where TRPC channels, TRP-1 and TRP-2, are mechanically activated by neck muscle contractions to influence neck steering behavior (Yeon et al., 2018), as well as the crab cardiac ganglion where mechanosensitive dendrites receive feedback from the postsynaptic muscles to modify the ganglion’s firing activity (García-Crescioni et al., 2010). Which receptors mediate VC mechanosensation is not known, but the VCs do express innexin gap junction proteins (Altun et al., 2009) that have recently been shown to act as mechanosensitive hemichannels (Walker & Schafer, 2020).

Serotonergic modulation of neural circuits and behavior is a well-characterized phenomena across both invertebrate and vertebrate animals (Bacqué-Cazenave et al., 2020; Weiger, 1997). In the mouse nociceptive circuit, serotonergic modulation has been shown to confer hyperalgesia (Bardoni, 2019; Lin et al., 2011). In this circuit, serotonin acts on dorsal root ganglion neurons to increase their sensitivity to mechanical stimuli through the 5-HT_2B_ receptor (Su et al., 2016), and to maintain this sensitivity through 5-HT_4/6/7_ receptors (Godínez-Chaparro et al., 2011). In the *C. elegans* egg-laying circuit, serotonin released by the HSN command neurons signals through several G-protein coupled receptors including the 5-HT_2_ ortholog SER-1 and the 5-HT_7_ ortholog SER-7 expressed on the vulval muscles (Hapiak et al., 2009; Hobson et al., 2006; Xiao et al., 2006). These serotonin receptors are thought to act through Ga_q_ and Ga_s_ signaling pathways, respectively, which activate EGL-19 L-type voltage-gated Ca^2+^ channels in the vulval muscles to enhance their response to other excitatory input (Schafer, 2006; Waggoner et al., 1998; Zhang et al., 2008). SER-7 is also expressed in the VC neurons (Fernandez et al., 2020), and serotonin acting through SER-7 has been shown to initiate motor neurons activity in other behaviors, such as feeding (Song et al., 2013). Like animals with blocked VC neurotransmission (Figure 1), *ser-7* mutants also fail to respond to exogenous serotonin (Hobson et al., 2006). Downstream targets of serotonergic signaling onto the VCs may include the N/P/Q-type Ca^2+^ channel UNC-2 to promote neurotransmitter release (Schafer et al., 1996). Following serotonergic potentiation, the VCs may be close enough to threshold to become mechanically excited in response to vm1-mediated vulval muscle twitches, leading to excitatory VC neurotransmitter release onto the vm2 vulval muscles to drive complete vulval muscle contraction for egg release.

Synaptic wiring diagrams show multiple sites of cross-innervation between the canonical egg-laying circuit and locomotion circuit (Cook et al., 2019; White et al., 1986). Consistent with this shared connectivity, egg-laying behavior is correlated with changes in locomotion behavior (Collins et al., 2016; Hardaker et al., 2001), suggesting that these two circuits actively communicate and coordinate their respective behaviors. Coordination of distinct neural circuits and behaviors has been demonstrated in the stomatogastric ganglion of the lobster, where two overlapping circuits are phase-coordinated with one another to regulate gastric peristalsis and subsequent digestion (Bartos et al., 1999; Clemens et al., 1998). The VCs have been shown to regulate locomotion, potentially through the GABAergic motor neurons or via direct release of ACh that drives excitation and contraction of the body wall muscles to slow locomotion (Figure8; Collins et al., 2016). Slowing of locomotion during egg laying may provide time for the vulval muscles to fully contract (Collins et al., 2016; Collins & Koelle, 2013). This slowing could function to hold the animal in a body posture that is favorable for vulval opening and egg release, which would be consistent with our finding that VC activity causes elongated vulval muscle Ca^2+^ twitch transients (Figure 3). In this role, the VCs would be signaling to the rest of the egg-laying circuit and locomotion circuit that the vulva is open so that body posture can be held in a favorable phase for egg release. The six VCs also make synapses onto each another (Cook et al., 2019). The mechanical activation of the vulva-proximal VCs during vulval opening may excite the distal VC neurons to slow locomotion and control body posture for efficient egg release. Failed egg-laying events could occur because of disrupted phasing of the locomotion pattern with activity in the egg-laying circuit. Since twitches and egg-laying events occur at a specific phase of the locomotion pattern (Collins & Koelle, 2013), the opening of the vulva and egg release may require not only coordinated muscle contraction, but also the relaxation of immediately proximal body wall muscles so that they cannot physically resist the full opening of the vulva. Thus, the VCs may act to coordinate the excitation of the vm2 muscles with excitatory input from the VA/VB locomotion motor neurons onto the vm1 muscles to facilitate full contraction and egg laying in phase with locomotion.

In all, the VC neurons and the egg-laying circuit present a tractable model system for investigating how different forms of chemical signaling and mechanosensory feedback work together to drive a robust behavior. The elucidation of the cellular and molecular mechanisms underlying these distinct forms of feedback could help the understanding of human neurological diseases where muscle coordination and proprioception are dysregulated, such as in Parkinson’s and Huntington disease (Bargmann, 2012; Lukos et al., 2013).

## Materials & Methods

### Nematode culture and strains

All *C. elegans* strains were maintained at 20 °C on Nematode Growth Medium (NGM) agar plates seeded with OP50 *E. coli* as described (Brenner, 1974). All assays were conducted on age-matched adult hermaphrodites at 24-30 h past the late L4 stage, unless otherwise stated. A list of all strains generated and used in this study can be found in Table 1.

**Table 1.** Strain names and genotypes for all animals used in this study.

| Strain | Genotype | Feature | Reference |
| --- | --- | --- | --- |
| N2 | *wild type* | Bristol wild-type strain | (Brenner, 1974) |
| CB190 | *unc-54(e190) I* | Myosin heavy chain null mutant | (Brenner, 1974) |
| LX1832 | *lite-1(ce314) X lin-15(n765ts) X* | Blue light-resistant (optogenetics and Ca<sup>2+</sup> imaging), multivulva (injection rescue marker) | (Collins & Koelle, 2013) |
| LX1918 | *vsIs164 lite-1(ce314) lin-15(n765ts) X* | Vulval muscles expressing GCaMP5, mCherry | (Collins et al., 2016) |
| LX1960 | *vsIs172; lite-1(ce314) lin-15(n765ts) X* | VC GCaMP5, mCherry | (Collins et al., 2016) |
| LX1970 | *wzIs30 IV; vsIs172; lite-1(ce314) lin-15(n765ts) X* | HSN Channelrhodopsin-2; VC GCaMP5, mCherry | (Collins et al., 2016) |
| LX1978 | *nlp-3(tm3023) X* | *nlp-3* null | (Brewer et al., 2019) |
| LX2004 | *vsIs183 lite-1(ce314) lin-15(n765ts) X* | HSN expressing GCaMP5, mCherry | (Collins et al., 2016) |
| MIA26 | *egl-1(n986dm) V* | No HSNs | (Ravi et al., 2018a) |
| MIA116 | *keyIs21; lite-1(ce314) lin-15(n765ts) X* | HSN HisCl | (Ravi et al., 2018a) |
| MIA123 | *egl-1(n986dm) V lite-1(ce314) lin-15(n765ts) X* | No HSNs | this study |
| MIA125 | *keyIs23; lite-1(ce314) lin-15(n765ts) X* | VC HisCl | this study |
| MIA144 | *keyIs33; lite-1(ce314) lin-15(n765ts) X* | VC TeTx | this study |
| MIA146 | *keyIs46; lite-1(ce314) lin-15(n765ts) X* | VC TeTx | this study |
| MIA173 | *keyIs46; egl-1(n986dm) V lite-1(ce314) lin-15(n765ts) X* | VC TeTx; no HSNs | this study |
| MIA176 | *keyIs23; egl-1(n986dm) V; lite-1(ce314) lin-15(n765ts) X* | VC HisCl; no HSNs | this study |
| MIA183 | *keyls33; vsIs164 lite-1(ce314) lin-15(n765ts) X* | VC TeTx; vulval muscle GCaMP5, mCherry | this study |
| MIA217 | *keyl33; vsIs183 lite-1(ce314) lin-15(n765ts) X* | VC TeTx; HSN GCaMP5, mCherry | this study |
| MIA221 | *keyls3; vsIs164; lite-1(ce314) lin-15(n765ts) X* | VC Channelrhodopsin-2; vulval muscle GCaMP5, mCherry | this study |
| MIA229 | *keyls48; lite-1(ce314) lin-15(n765ts) X* | Vulval muscle Channelrhodopsin-2 | this study |
| MIA241 | *keyls48; vsIs172; lite-1(ce314) lin-15(n765ts) X* | Vulval muscle Channelrhodopsin-2; VC GCaMP5, mCherry | this study |
| MIA250 | *keyls48; vsIs164 lite-1(ce314) lin-15(n765ts) X* | Vulval muscle Channelrhodopsin-2, GCaMP5, mCherry | this study |
| MIA298 | *keyls48; vsIs172; unc-54(e190) I; lite-1(ce314) lin-15(n765ts) X* | Vulval muscle Channelrhodopsin-2; VC GCaMP5, mCherry; *unc-54* myosin heavy chain null mutant background | this study |
| MIA428 | *nlp-3(tm3023); keyls33* | VC TeTx; *nlp-3* null | this study |

### Plasmid and strain construction

Oligonucleotides were synthesized by IDT-DNA. PCR was performed using high-fidelity Phusion DNA polymerase (New England Biolabs) except for routine genotyping which was performed using standard Taq DNA polymerase. Plasmids were prepared using a Qiagen miniprep spin kit. DNA concentrations were determined using a Nano-Drop spectrophotometer.

#### Tetanus toxin (TeTx)-expressing transgenes

##### VC neuron TeTx

A ∼1.4 kB DNA fragment encoding TeTx was cut from pAJ49 (Jose et al., 2007) with AgeI/XhoI, and ligated into a similarly digested pKMC145 [*lin-11::GFP::unc-54 3’ UTR*] to generate pKMC282 [*lin-11::TeTx::unc-54 3’ UTR*]. pKMC282 (50 ng/µl) was injected along with pL15EK (50 ng/µl; (Clark et al., 1994)) into LX1832 *lite-1(ce314) lin-15(n765ts) X* generating four independent transgenic lines of which one, MIA113 *keyEx32 [lin-11::TeTx::unc-54 3’UTR* + *lin-15(+)]; lite-1(ce314) lin-15(n765ts) X*, was used for integration. *keyEx32* was integrated with UV/TMP generating three independent integrants *keyIs32-33* and *keyIs46 [lin-11::TeTx::unc-54 3’UTR* + *lin-15(+)]*. Each transgenic line was backcrossed six times to LX1832 generating strains MIA144-146. All transgenic strains appeared phenotypically similar, and MIA144 and MIA146 were used for experiments and further crosses. To eliminate HSNs in animals lacking VC synaptic transmission, MIA26 *egl-1(n986dm) V* mutant animals were crossed with MIA146 to generate MIA173 *keyIs46; egl-1(n986dm) V*; *lite-1(ce314) lin-15(n765ts) X*.

#### *Histamine-gated chloride channel (HisCl)-expressing transgenes*

##### VC neuron HisCl

The ∼1.3 kB DNA fragment encoding *Drosophila* histamine-gated chloride channel HisCl1 was PCR amplified from pNP403 (Pokala et al., 2014) using the oligonucleotides 5’-GCG CCC GGG GTA GAA AAA ATG CAA AGC CCA ACT AGC AAA TTG G-3’ and 5’-GCG GAG CTC TTA TCA TAG GAA CGT TGT CCA ATA GAC AAT A-3’, cut with XmaI/SacI, and ligated into AgeI/SacI-digested pKMC145 to generate pAB2 [*lin-11::HisCl::unc-54 3’UTR]*. pAB2 (20 ng/µl) was injected into LX1832 along with pL15EK (50 ng/µl), generating the extrachromosomal line MIA93 *keyEx24 [lin-11::HisCl::unc-54 3’UTR* + *lin-15(+)]; lite-1(ce314) lin-15(n765ts) X*. The extrachromosomal transgene subsequently integrated using UV/TMP to generate the transgenes *keyIs23*-30 *[lin-11::HisCl::unc-54 3’UTR* + *lin-15(+)]*. Strains bearing these transgenes were then backcrossed to the LX1832 parent strain six times, generating strains MIA124, MIA125, MIA130, MIA131, and MIA132. All transgenic strains appeared phenotypically similar, and MIA125 was used for experiments and further crosses. To eliminate the HSN neurons in animals expressing HisCl in the VC neurons, MIA125 *keyIs23; lite-1(ce314) lin-15(n765ts) X* was crossed with MIA26 to generate MIA176 *keyIs23; egl-1(n986dm)*; *lite-1(ce314) lin-15(n765ts) X*.

#### Channelrhodopsin-2 (ChR2)-expressing transgenes

##### Vulval muscle ChR2

To express ChR2 in the vulval muscles, the ∼1 kB DNA fragment encoding for ChR2 was PCR amplified from pRK7 [*del-1::ChR2(H34R/T159C::unc-54 3’ UTR]* using oligonucleotides 5’-GCG GCT AGC ATG GAT TAT GGA GGC GCC CTG-3’ and 5’-GCG GGT ACC TCA GGT GGC CGC GGG GAC CGC GCC AGC CTC GGC C-3’. The amplicon and recipient plasmid, pBR3 (Ravi et al., 2018a), were digested with NheI/KpnI, generating pRK11 [*ceh-24::ChR2(H34R/T159C)::unc-54 3’ UTR*]. pRK11 (50 ng/µl) was injected into LX1832 along with pL15EK (50 ng/µl) generating MIA212 *keyEx43* [*ceh-24::ChR2(H34R/T159C)::unc-54 3’UTR + lin-15(+)]* which was subsequently integrated with UV/TMP, generating five independent integrated transgenes *keyIs47-51*. Strains carrying these integrated transgenes were then backcrossed to the LX1832 parent strain six times, generating the strains MIA229-232 and MIA242. All transgenic strains were phenotypically similar, and MIA229 was used for experiments and further crosses.

##### VC ChR2

The allele *keyIs3* was used to express ChR2 in the VC neurons under a modified *lin-11* promoter, as previously described (Collins et al., 2016).

##### HSN ChR2

The allele *wzIs30* was used to express ChR2 in the HSN neurons under the *egl-6* promoter, as previously described (Collins et al., 2016; Emtage et al., 2012).

#### Calcium reporter transgenes

##### Vulval muscle GCaMP5

Vulval muscle Ca^2+^ activity was visualized using the strain LX1918 which co-expresses GCaMP5 and mCherry in the vulval muscles from the *unc-103e* promoter (Collins et al., 2016). To analyze vulval muscle Ca^2+^ activity in animals where VC synaptic transmission was blocked with TeTx, LX1918 was crossed with MIA144 to generate MIA183 *keyIs33*; *vsIs164 lite-1(ce314) lin-15(n765ts) X*. To analyze vulval muscle Ca^2+^ activity in animals where the VC neurons could be optogenetically activated by ChR2, LX1918 was crossed with MIA3 (Collins et al., 2016), to generate MIA221 *keyIs3; vsIs164 lite-1(ce314) lin-15(n765ts) X*. To analyze vulval muscle Ca^2+^ activity in animals where the vulval muscles could be optogenetically activated by ChR2, LX1918 was crossed with MIA229 to generate MIA250 *keyIs49*; *vsIs164 lite-1(ce314) lin-15(n765ts) X*.

##### VC neuron GCaMP5

VC neuron Ca^2+^ activity was visualized using the strain LX1960 which co-expresses GCaMP5 and mCherry in the VC neurons (Collins et al., 2016). To visualize VC activity during optogenetic stimulation of the HSNs, the strain LX1970 was used (Collins et al., 2016). To visualize VC activity after optogenetic stimulation of the vulval muscles, LX1960 was crossed with MIA229 to generate MIA241 *vsIs172*; *keyIs48 lite-1(ce314) lin-15(n765ts) X*. To visualize VC activity after optogenetic stimulation of the vulval muscles when muscle contraction is impaired, CB190 *unc-54(e190) I* myosin heavy chain null mutants were crossed with LX1832 to generate MIA274 *unc-54(e190) I*; *lite-1(ce314) lin-15(n765ts) X*. MIA274 was then crossed with MIA241 to generate MIA298 *keyIs48; vsIs172; unc-54(e190) I; lite-1(ce314) lin-15(n765ts) X*.

##### HSN neuron GCaMP5

HSN neuron Ca^2+^ activity was visualized using the strain LX2004 which co-expresses GCaMP5 and mCherry in the HSN neurons (Collins et al., 2016). To visualize HSN Ca^2+^ after neurotransmission from the VC neurons is blocked by TeTx, LX2004 was crossed with MIA144 to generate MIA217 *keyIs33; vsIs183 lite-1(ce314) lin-15(n765ts) X*.

### Ratiometric Ca^2+^ imaging

Ratiometric Ca^2+^ imaging of the vulval muscles and VC neurons in freely behaving animals was performed as previously described methods (Ravi et al., 2018b). Late L4 hermaphrodites were staged and then imaged 24 h later by being moved to an NGM agar chunk between two glass coverslips. Animals were recorded on a Zeiss Axio Observer.Z1 inverted compound microscope with a 20X 0.8NA Apochromat objective. Brightfield recordings of behavior was recorded with infrared illumination using a FLIR Grasshopper 3 CMOS camera after 2×2 binning using FlyCap software. Colibri.2 470 nm and 590 nm LEDs were used to co-excite GCaMP5 and mCherry fluorescence which was captured at 20 Hz onto a Hamamatsu ORCA Flash-4.0V2 sCMOS camera after channel separation using a Gemini image splitter. For Ca^2+^ imaging of the vulval muscles, a lateral focal plane was used to capture the anterior and posterior vm1 and vm2 cells on one side of the animal. For Ca^2+^ imaging of the VC neurons, a lateral focal plane was used to capture the VC4 and VC5 cell bodies and their presynaptic termini around the vulva. Each animal was recorded until it entered an active egg-laying phase (up to 1 h), after which a 10-minute segment centered around the onset of egg laying was extracted from the full recoding for analysis. Two-channel fluorescence (GCaMP5/mCherry) image sequences were processed and analyzed in Fiji (Schindelin et al., 2012), Volocity (PerkinElmer), and a custom script for MATLAB (MathWorks) as previously described (Ravi et al., 2018b).

Ratiometric Ca^2+^ imaging of the HSN neurons in freely behaving animals was performed as previously described (Ravi et al., 2018a). Late L4 hermaphrodites were staged and then imaged 24 h later by being moved to an NGM agar chunk and a glass coverslip being placed over. Animals were recorded on an inverted Leica TCS SP5 confocal microscope with a 20X 0.7NA Apochromat objective. 488 nm and 561 nm laser lines were used to co-excite GCaMP5 and mCherry fluorescence, respectively.

### Electrical silencing using HisCl

Acute electrical silencing with histamine was performed as previously described (Pokala et al., 2014; Ravi et al., 2018a). Animals were moved onto OP50-seeded NGM agar plates that contained either 0 mM or 10 mM histamine for four hours before the experiment.

### Egg-laying behavior assays

The steady-state accumulation of eggs in the uterus was determined as previously described (Koelle & Horvitz, 1996). Briefly, late L4 hermaphrodites were staged onto OP50-seeded plates and grown at 20 °C for 30 h after which a single adult was placed into a 7 ul drop of 20% sodium hypochlorite (bleach) solution. The eggs, which are resistant to bleach, were then counted using a dissecting microscope. The timing of the first egg laid was assayed by staging a young adult (within 30 minutes of L4 to adult molt) animal onto an NGM agar plate with food and checking every following 30 minutes for the presence of an egg on the plate (Ravi et al., 2018a).

Egg laying in liquid in response to exogenous serotonin was performed as described (Banerjee et al., 2017; Trent et al., 1983). Late L4 hermaphrodites were staged onto OP50-seed plates and grown at 20 °C for 24 h. Adult animals were placed singly into either 100 ul M9 buffer only or M9 buffer containing 18.5 mM serotonin creatinine sulfate salt (Sigma-Aldrich) in a 96-well microtiter dish. The number of eggs laid by each animal after 1 hour were then counted. Egg laying on plates in response to exogenous serotonin was performed on NGM agar infused with

18.5 mM serotonin creatine sulfate salt (Sigma-Aldrich). 3 animals were placed on each NGM agar plate and the number of eggs laid was counted after 1 hour and divided by 3 to calculate an average-animal response per plate.

### Optogenetics

Optogenetic experiments with Channelrhodopsin-2 (ChR2) were performed using a Zeiss Axio Observer.Z1 inverted compound microscope as previously described (Collins et al., 2016). ChR2 was excited in freely behaving animals using a 470 nm (blue) LED. Late L4 animals were staged onto NGM agar plates seeded with *E. coli* OP50 bacterial cultures containing either 0.4 mM all-*trans* retinal (ATR) or no ATR 24 h prior to the start of the experiment. Animals were then continuously illuminated with blue light for 3-5 minutes and the locomotion behavior and number of egg-laying events were recorded. For experiments combining optogenetics with ratiometric Ca^2+^ imaging, the blue light would excite both the ChR2 and GCaMP5 fluorescence simultaneously. Blue light intensity was chosen based on both optimal settings to observe robust ChR2-activation and GCaMP5 fluorescence. Animals were excluded from a dataset if they entered an active egg-laying state before the onset of blue light stimulation (one animal in total across all experiments).

### Experimental design and statistical analyses

Sample sizes for behavioral assays and Ca^2+^ imaging experiments followed previous studies (Banerjee et al., 2017; Collins et al., 2016; Ravi et al., 2018a). A minimum of 10 animals per genotype per condition were measured across all experiments. One animal was excluded from the analysis of VC Channelrhodopsin-2; vulval muscles GCaMP5 experiment because it entered an egg-laying active state before the onset of blue light stimulation. No other data or animals were excluded. No explicit power analysis was performed prior to the study. All data were analyzed using GraphPad Prism 8. Steady-state egg accumulation and timing of first egg laid assays were compared using a one-way ANOVA with Bonferroni’s correction for multiple comparisons. Serotonin-induced egg laying assays were compared using either a Kruskal-Wallis test with Dunn’s correction for multiple comparisons (in buffer with individual responses) or a one-way ANOVA with Bonferroni’s correction for multiple comparisons (on plates with averaged responses). Inter-transient intervals of Ca^2+^ transients compared between active and inactive egg-laying behavior states were pooled together across all animals and analyzed using either a Kolmogorov-Smirnov test or Kruskal-Wallis test with Dunn’s correction for multiple comparisons. Per animal rates of failed egg-laying events were analyzed using a Mann-Whitney test. Ca^2+^ transient peak amplitude, inter-transient interval, or transient widths were first averaged across each animal and then these averages were compared across animals using either a Student’s t test or a one-way ANOVA with Bonferroni’s correction for multiple comparisons. The number of animals and instances of analyzed behavior events along with exact p values resulting from defined statistical tests are reported within each figure legend.

## Acknowledgements

This work was funded by grants from the NIH (R01-NS086932) and NSF (IOS-1844657) to KMC. RJK 3^rd^ was supported by a University of Miami Maytag Fellowship. We thank David M. Miller III for sharing plasmids. Some of the strains used in this study were provided by the *C. elegans* Genetics Center, which is funded by NIH Office of Research Infrastructure Programs (P40 OD010440). We thank Drs. Julia Dallman, Grace Zhai, Mason Klein, and members of the Collins lab for helpful discussions and feedback on the manuscript.

---

**Movie 1. GCaMP5/mCherry ratio recording of VC Ca^2+^ activity during the egg-laying active state**

Ratio of GCaMP5 and mCherry fluorescence in the VC neurons mapped onto a false color spectrum ranging from blue (low Ca^2+^) to red (high Ca^2+^). The VC neurons show high Ca^2+^ activity and physical displacement as the vulval muscles contract and an egg passes through the vulva to be laid. The Ca^2+^ activity then returns to a low level until a vulval muscle twitch occurs (0:09) followed by another egg-laying event.

**Movie 2. GCaMP5/mCherry ratio recording of vulval muscle Ca^2+^ in response to optogenetic stimulation of the VCs**

Ratio of GCaMP5 and mCherry fluorescence in the vulval muscles mapped onto a false color spectrum ranging from blue (low Ca^2+^) to red (high Ca^2+^). Optogenetic activation of the VC neurons induces vulval muscle Ca^2+^ activity and twitches, but not egg laying.

**Movie 3. GCaMP5/mCherry ratio recording of vulval muscle Ca^2+^ during the egg-laying active state when synaptic transmission from the VC neurons is blocked by TeTx**

Ratio of GCaMP5 and mCherry fluorescence in the vulval muscles mapped onto a false color spectrum ranging from blue (low Ca^2+^) to red (high Ca^2+^). The vulval muscles still exhibit twitches and egg-laying events but will also frequently fail to lay eggs in response to high Ca^2+^ activity, termed “failed egg-laying events”.

**Movie 4. Brightfield recording of vulval opening and egg laying in response to optogenetic stimulation of the vulval muscles**

Brightfield recording of egg laying in response to optogenetic vulval muscle activation. Optogenetic activation of the vulval muscles induces tetanic vulval muscle contraction, but vulval opening and egg release remains phased with the body bends of locomotion.

**Movie 5. GCaMP5/mCherry ratio recording of VC Ca^2+^ in response to optogenetic stimulation of the vulval muscles**

Ratio of GCaMP5 and mCherry fluorescence in the VCs mapped onto a false color spectrum ranging from blue (low) to red (high). Optogenetic activation of the vulval muscles causes an immediate induction of VC Ca^2+^ activity that remains at a high level for the duration of the stimulation.

